## Supplemental Information for "Tracking N- and C-termini of *C. elegans* polycystin-1 reveals their distinct targeting requirements and functions in cilia and extracellular vesicles"

Supplementary Information:

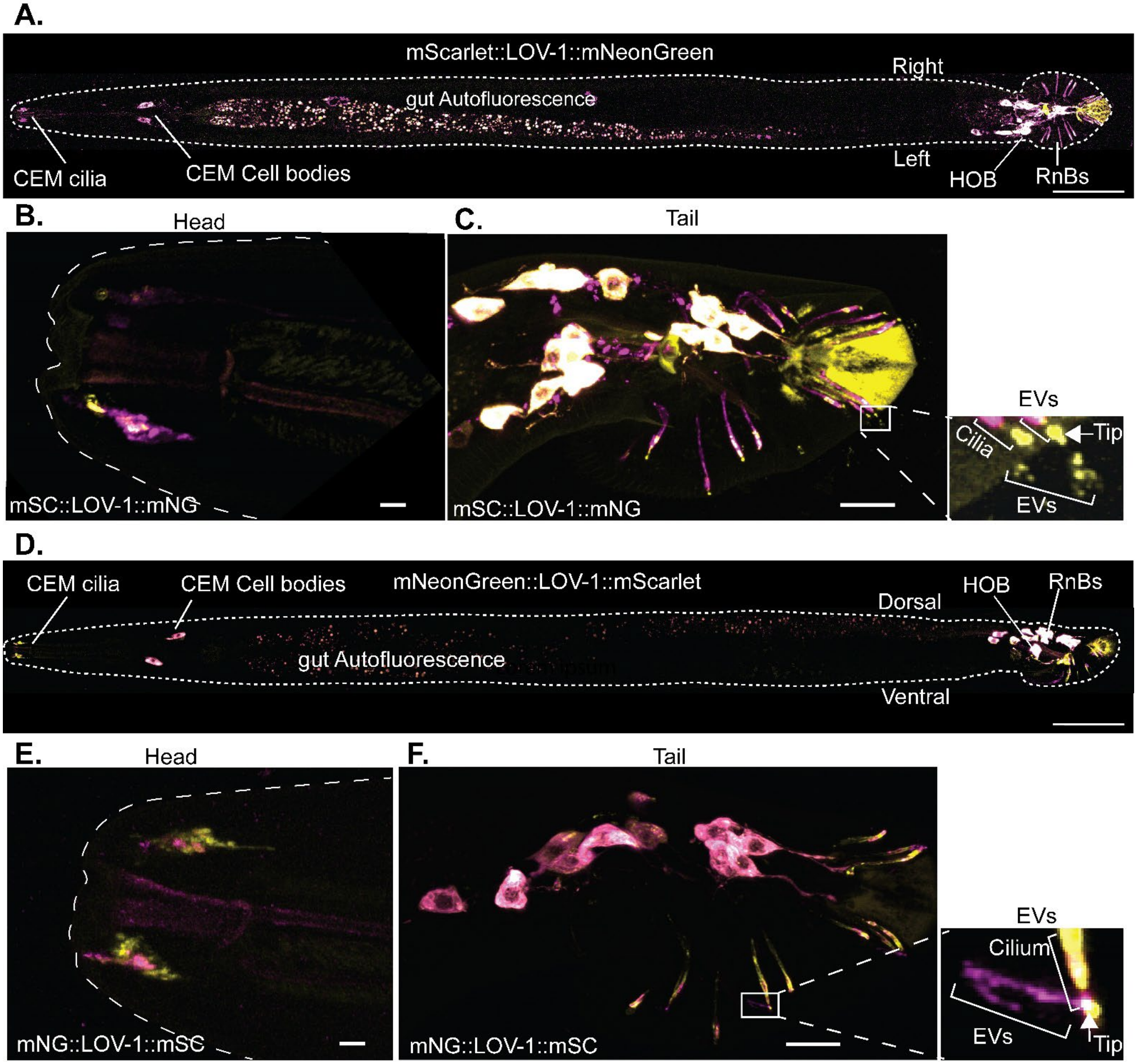

<u>S1 Fig.</u> **CRISPR-tagged LOV-1 reporters have similar localization patterns when tagged with different fluorescent proteins at each terminus**. A) 40X tile scan of a whole *C. elegans* male expressing mSC::LOV-1::mNG. Scale bar is 50 uM. B) Z-projection of male head of worm expressing mSC::LOV-1::mNG. C) Z-projection of male tail of worm expressing mSC::LOV-1::mNG. Inset shows CTM LOV-1 (LOV-1::mNG) EVs released from the cilia tips of RnB neuronal cilia. D) 40X tile scan of a whole male worm expressing mNG::LOV-1:: mSC. Scale bar is 50 uM. E) Z-projection of male head of worm expressing mNG::LOV-1::mSC. F) Z-projection of male tail of worm expressing mNG::LOV-1::mSC. Inset shows CTM LOV-1 (LOV-1::mSC) EVs released from the cilia tips of RnB neuronal cilia. The signal in the green channel at the ciliary tip is autofluorescence of the cuticular pore. If we reduce intensity,

autofluorescent signal is absent but EVs cannot be imaged. B, C, E, F) Scale bars for head and tail images are 2um and 10um respectively.

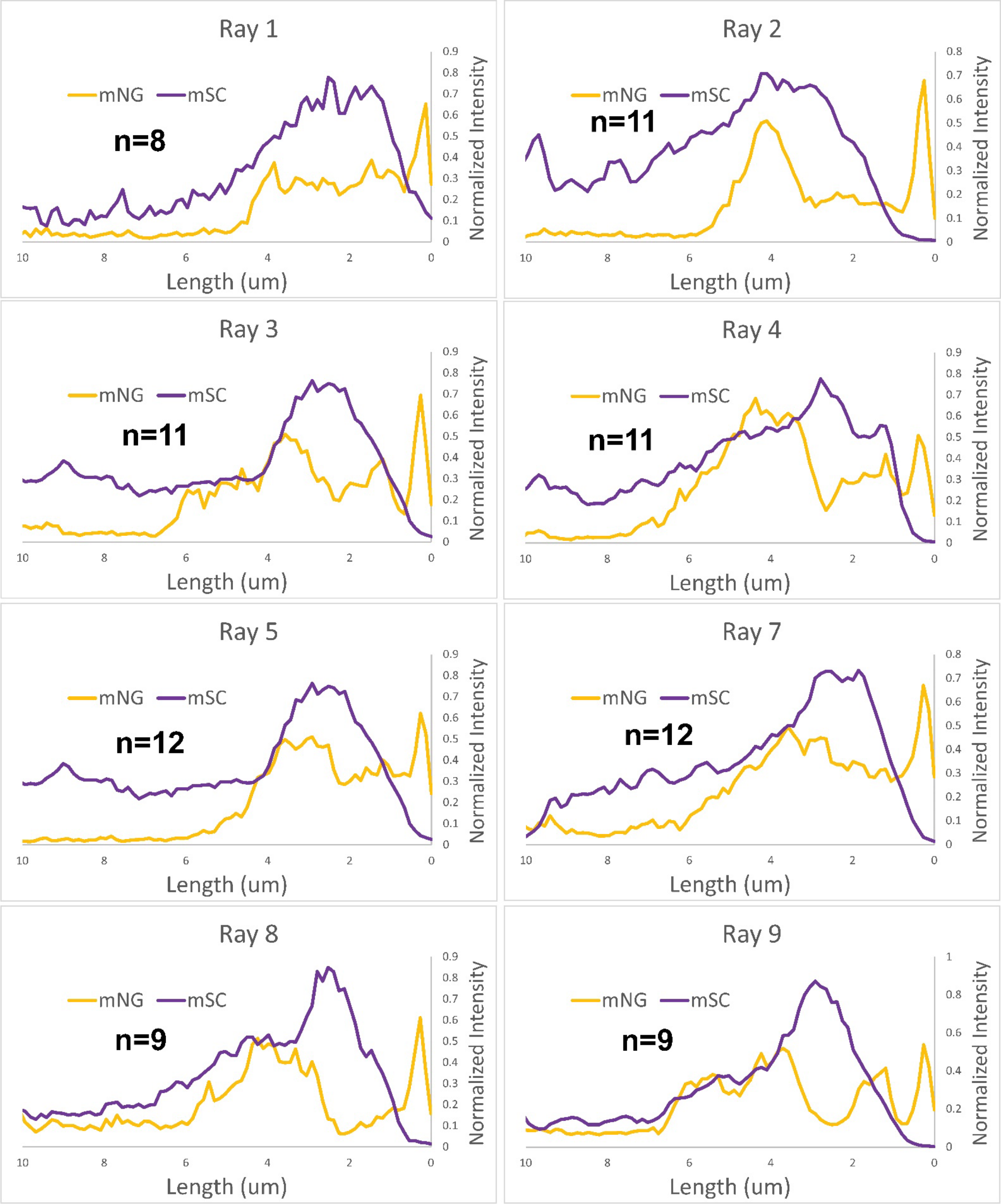

**S2 Fig. All eight polycystin expressing ray (RnB) neurons show the same pattern of NTM (mSC) and CTM (mNG) enrichment at the distal dendrites and cilia when expressing mSC::LOV-1::mNG.** NTM LOV-1 (mSC::LOV-1) is enriched at the distal dendrite, cilia base, and cilia proper but is absent from cilia tips. CTM LOV-1 (LOV-1::mNG) is enriched at the cilia base and cilia tip but not the distal dendrite. n= number of ray neurons measured.

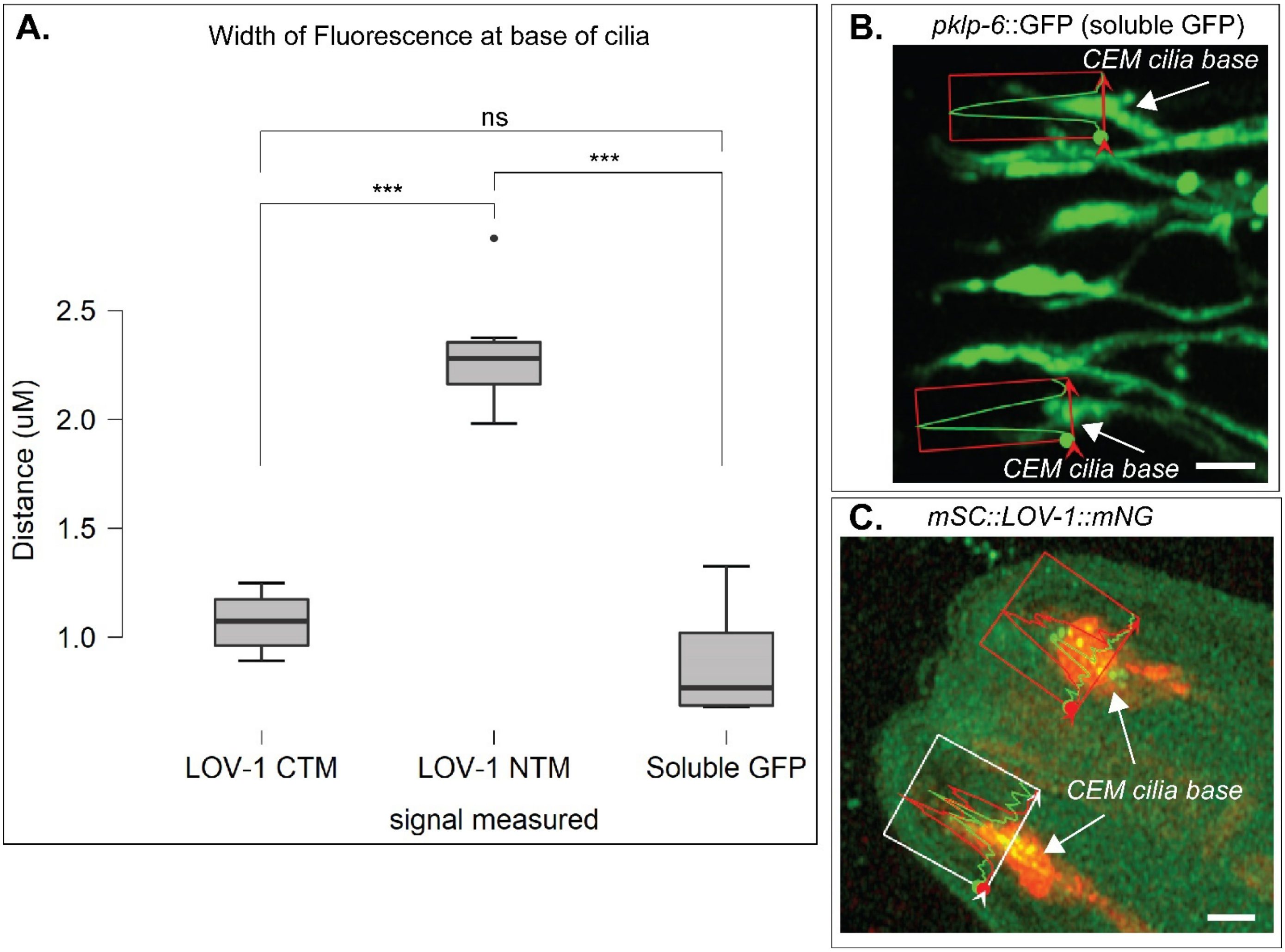

**<u>S3 Fig.</u> NTM LOV-1 is released outside the CEM cilium.** A) comparison of the width of fluorescent signal at the base of cilia in mSC::LOV-1::mNG worms compared to a soluble GFP ciliated neuronal marker. Statistics performed was an ANOVA with Tukey post-hoc analysis for multiple comparisons. B) representative image of *pklp-6*::GFP in the male head. CEM and IL2 cilia are visible. Width measurements were limited to the CEM cilia base region. C) representative image of mSC::LOV-1::mNG used for signal width measurements. LOV-1 CTM and CEM-expressed soluble GFP correlate in width, thus NTM must be released outside the CEM cilium. N=6, 6, and 5 for LOV-1 CTM, LOV-1 NTM, and soluble GFP respectively. Statistics performed using one-way ANOVA and Tukey's post hoc test. Scale bars are 2uM.

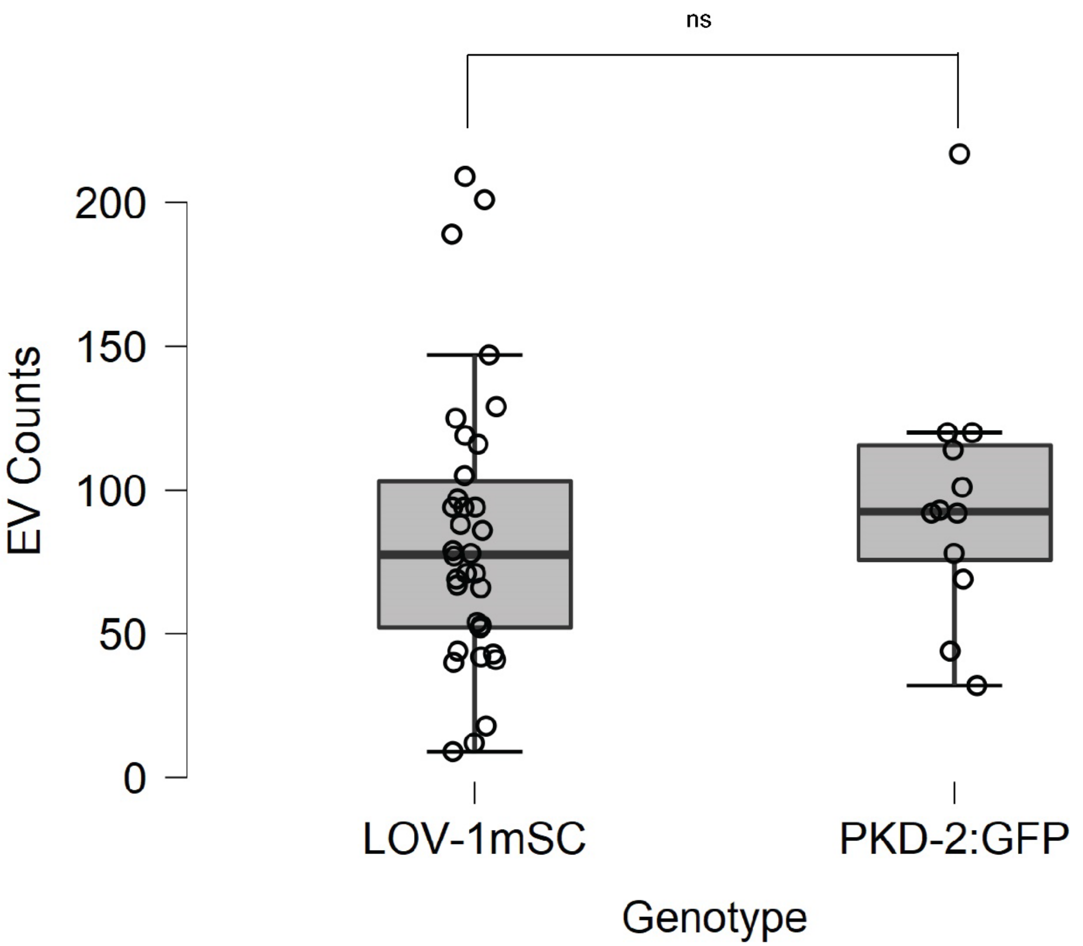

**Descriptive Statistics**

| | EV Counts | |
|---|---|---|
| | **LOV-1mSC** | **PKD-2:GFP** |
| N | 34 | 12 |
| Mean | 84.676 | 97.667 |
| Std. Deviation | 49.034 | 46.713 |
| Minimum | 9.000 | 32.000 |
| Maximum | 209.000 | 217.000 |

**Independent Samples T-Test**

| | t | df | p |
|---|---|---|---|
| EV Counts | -0.817 | 20.201 | 0.423 |

*Note.* Welch's t-test.

**S4 Fig. Endogenous LOV-1::mSC and transgenic PKD-2 are released from male tail in EVs outside the worm at the same abundance in single expression strains.** Slightly different EV counts are due to background and/or EV movement during imaging. EV counts were done using an automated counting plugin, ComDet, in ImageJ. Stats generated in JASP and Welch’s T-test was performed.

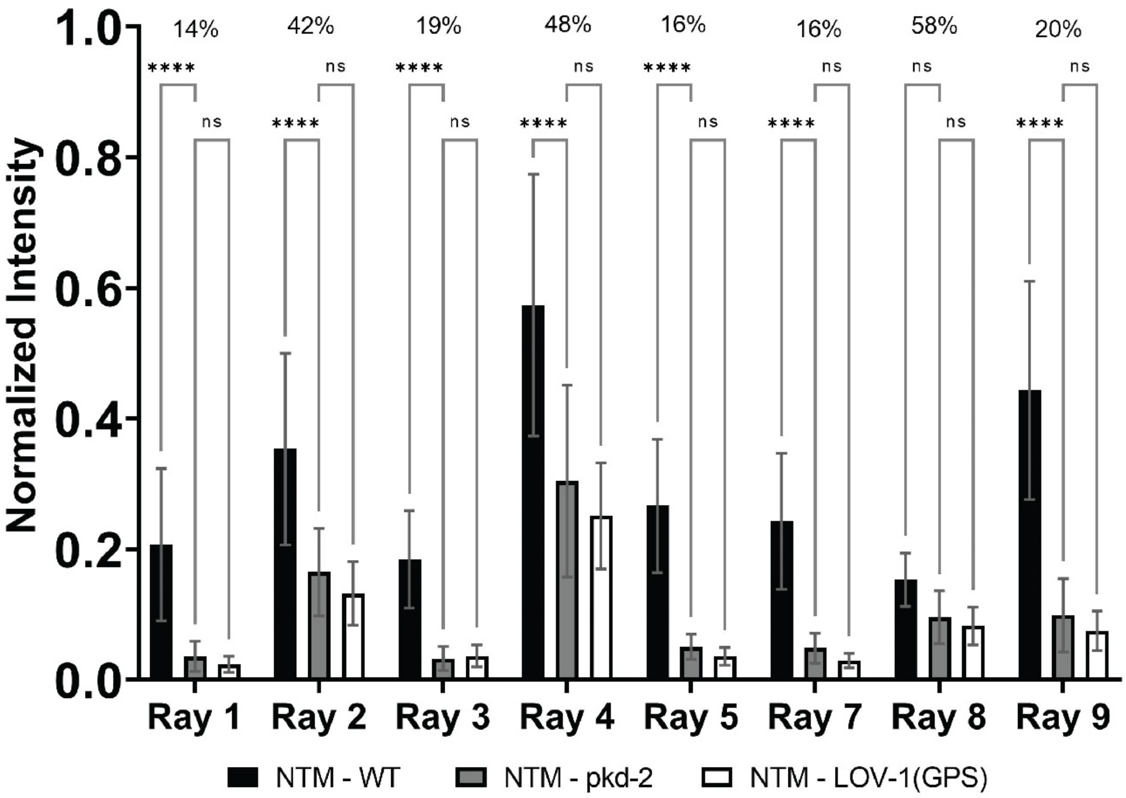

**S5 Fig. *pkd-2* and LOV-1 GPS are required for proper localization of NTM LOV-1 to ray dendrites and cilia.** Fluorescence profiling of NTM LOV-1 (mScarlet) in ray cilia of worms expressing mSC::LOV-1::mNG in WT, *pkd-2(sy606),* or *msc::lov-1(c2181s)::mng* backgrounds. Percentages above the graph represent the percent reduction of signal in the mutant strains compared to WT. Rays 2, 4, and 8 (the ventral rays) show an enrichment in the mutant strains when compared to the other rays. Statistics performed using 2-way ANOVA and Tukey's multiple comparisons tests. Sample sizes provided in source data.

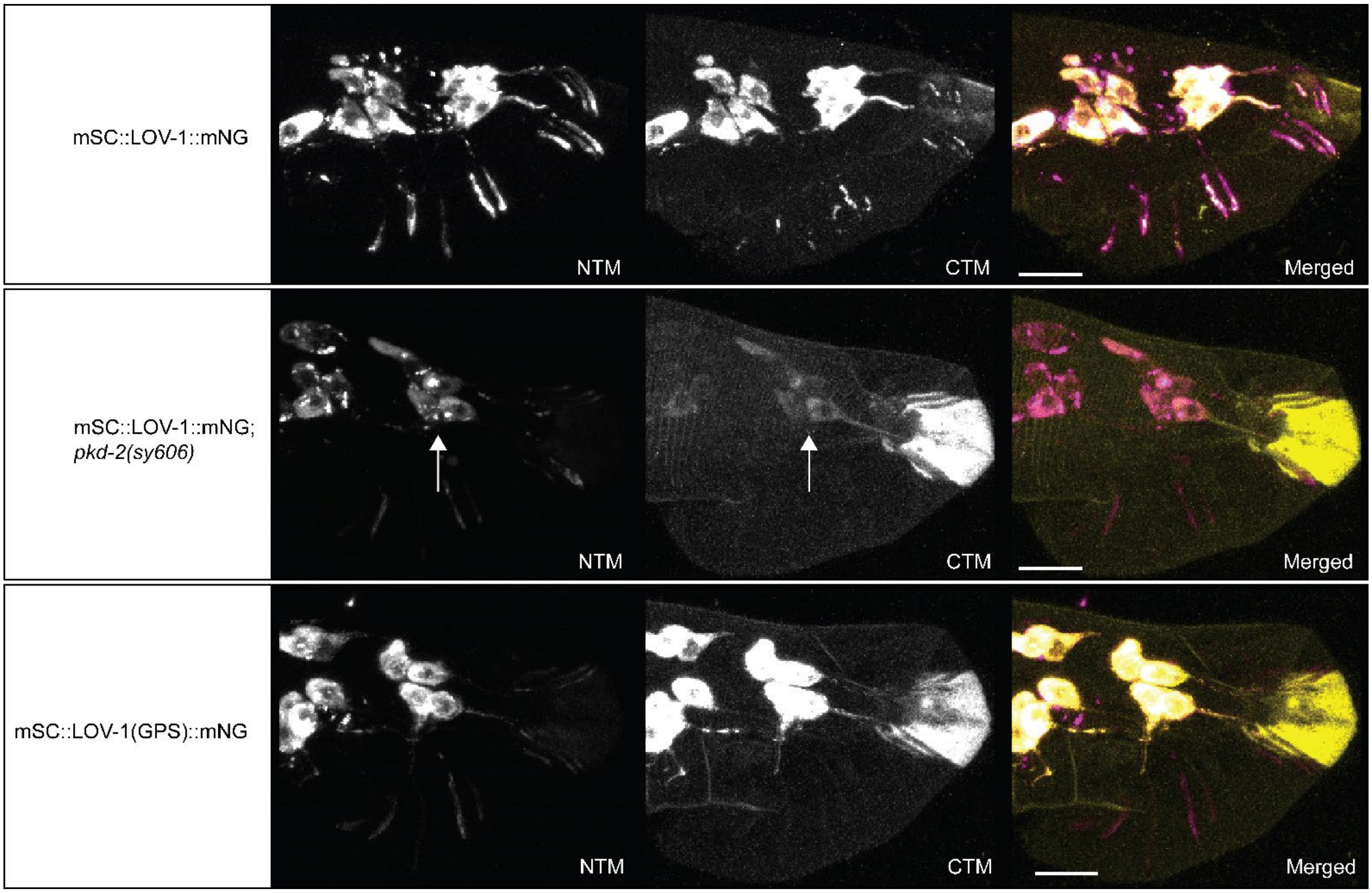

**S6 Fig. Cell body abundance of CTM LOV-1 is reduced in *pkd-2(sy606)* when compared to WT and *lov-1(GPS)* strains (arrows).**

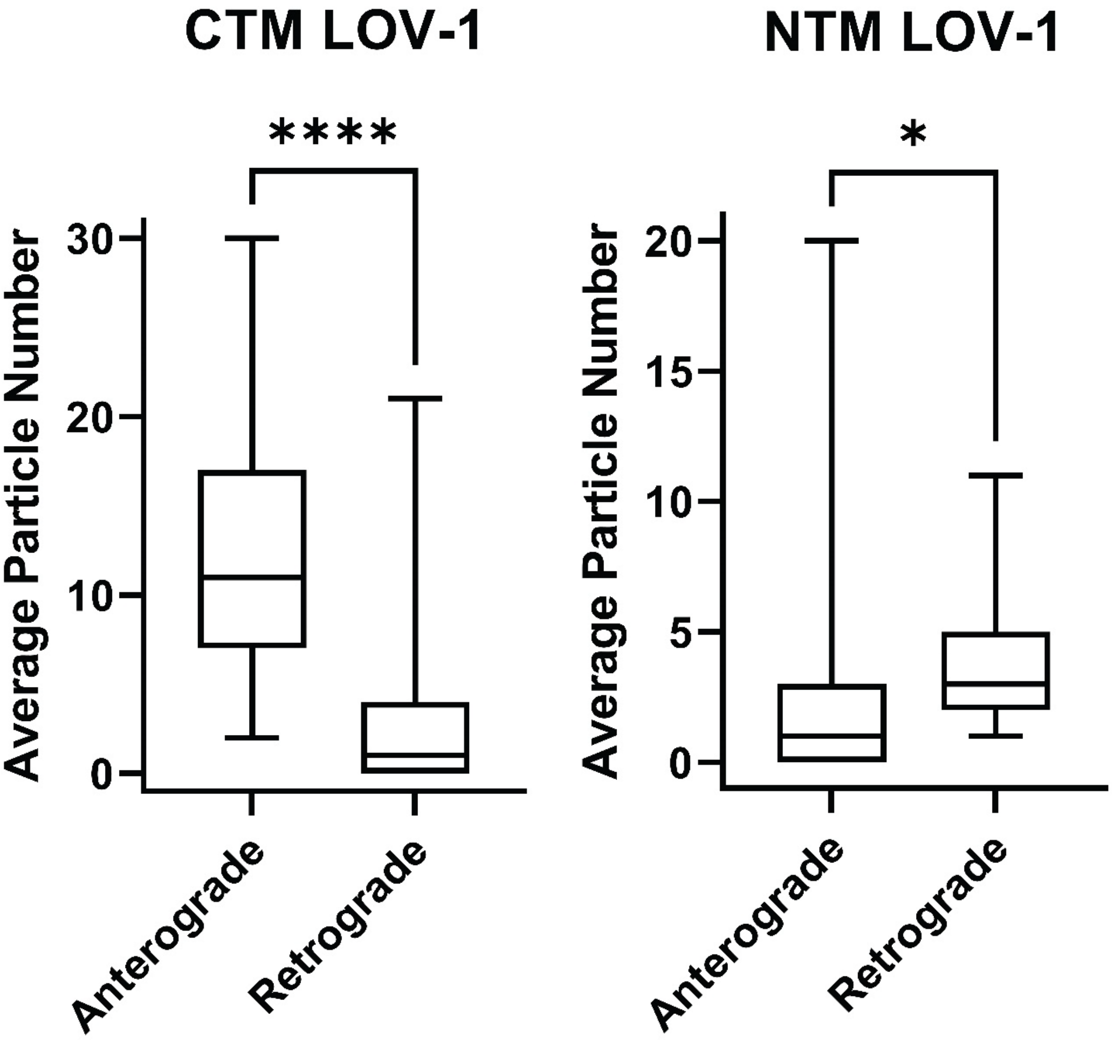

**<u>S7 Fig.</u> Transport event per kymograph.** CTM LOV-1 transport occurs more frequently in the anterograde direction, whereas NTM LOV-1 transport occurs more frequently in the retrograde direction in dendrites. N = 46 and 56 for CTM LOV-1 and NTM LOV-1 respectively. Statistics performed was unpaired t-test.

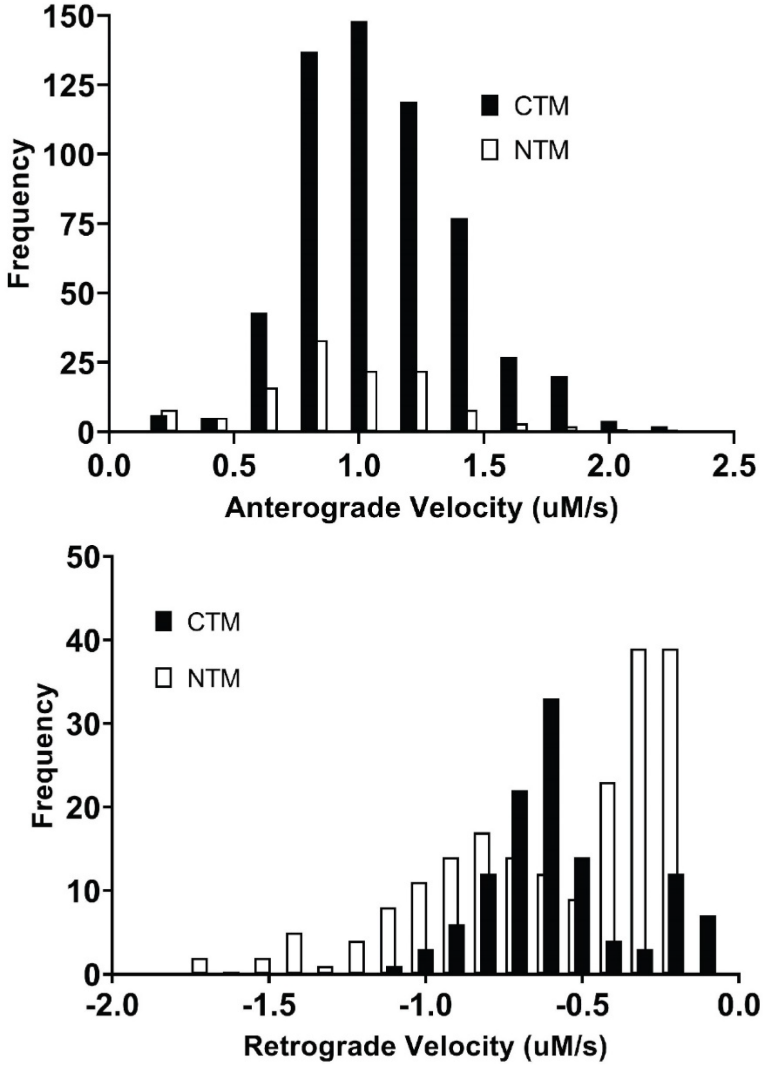

| Descriptive Statistics | | | | |
|---|---|---|---|---|
| | Velocity | | | |
| | CTM/Anterograde | CTM/Retrograde | NTM/Anterograde | NTM/Retrograde |
| N | 588 | 117 | 120 | 200 |
| Mean | 1.066 | -0.568 | 0.911 | -0.479 |
| Std. Deviation | 0.32 | 0.227 | 0.357 | 0.36 |
| Minimum | 0.139 | -1.05 | 0.143 | -1.621 |
| Maximum | 2.263 | -0.091 | 1.946 | -0.052 |

**<u>S8 Fig.</u> Histograms showing the velocity distributions of CTM and NTM LOV-1 in anterograde and retrograde directions in dendrites.**

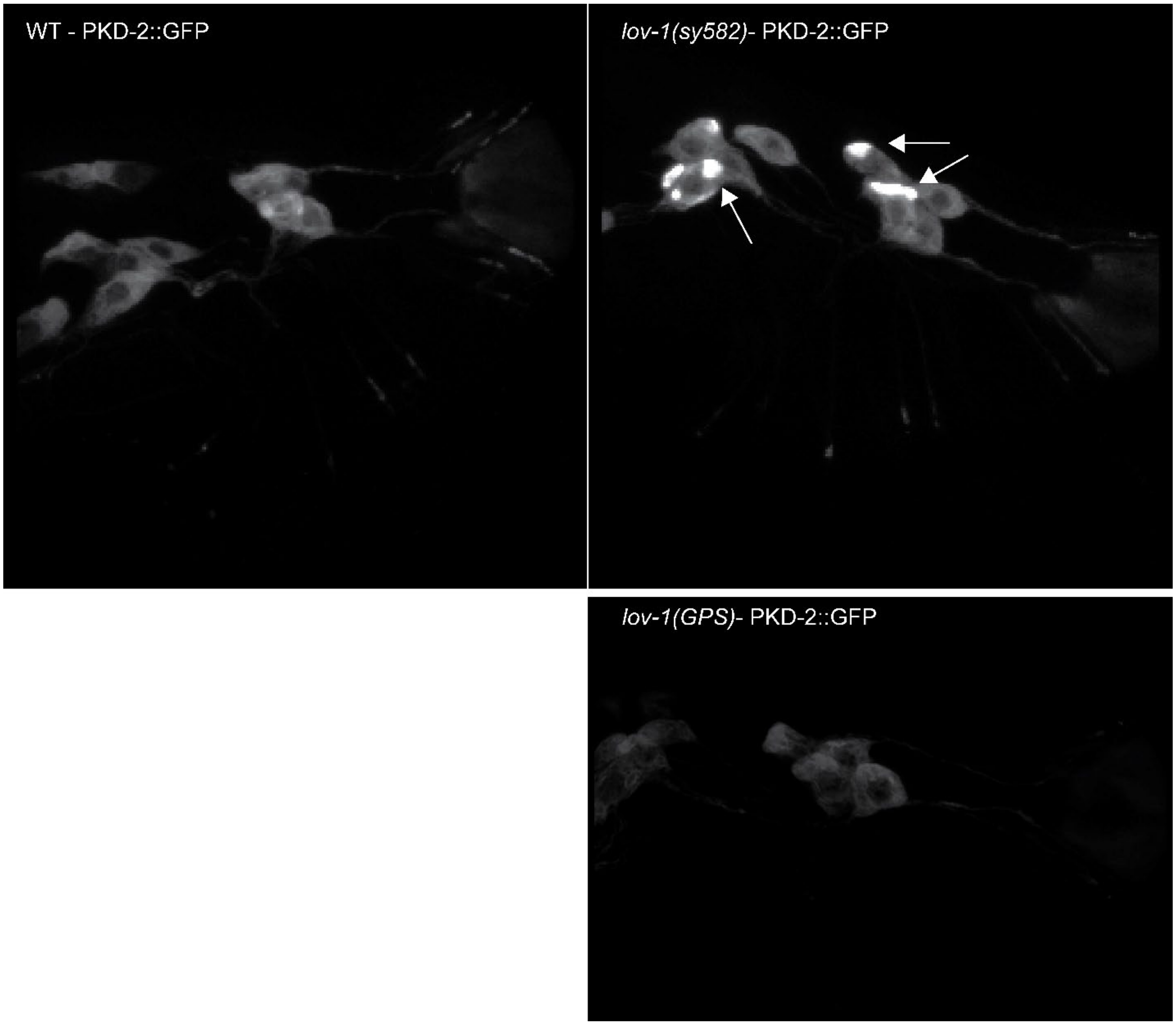

**<u>S9 Fig.</u> PKD-2 accumulation in cell bodies of ray neurons is observed in *lov-1(sy582)* deletion mutants but not *lov-1(GPS)* mutants**.

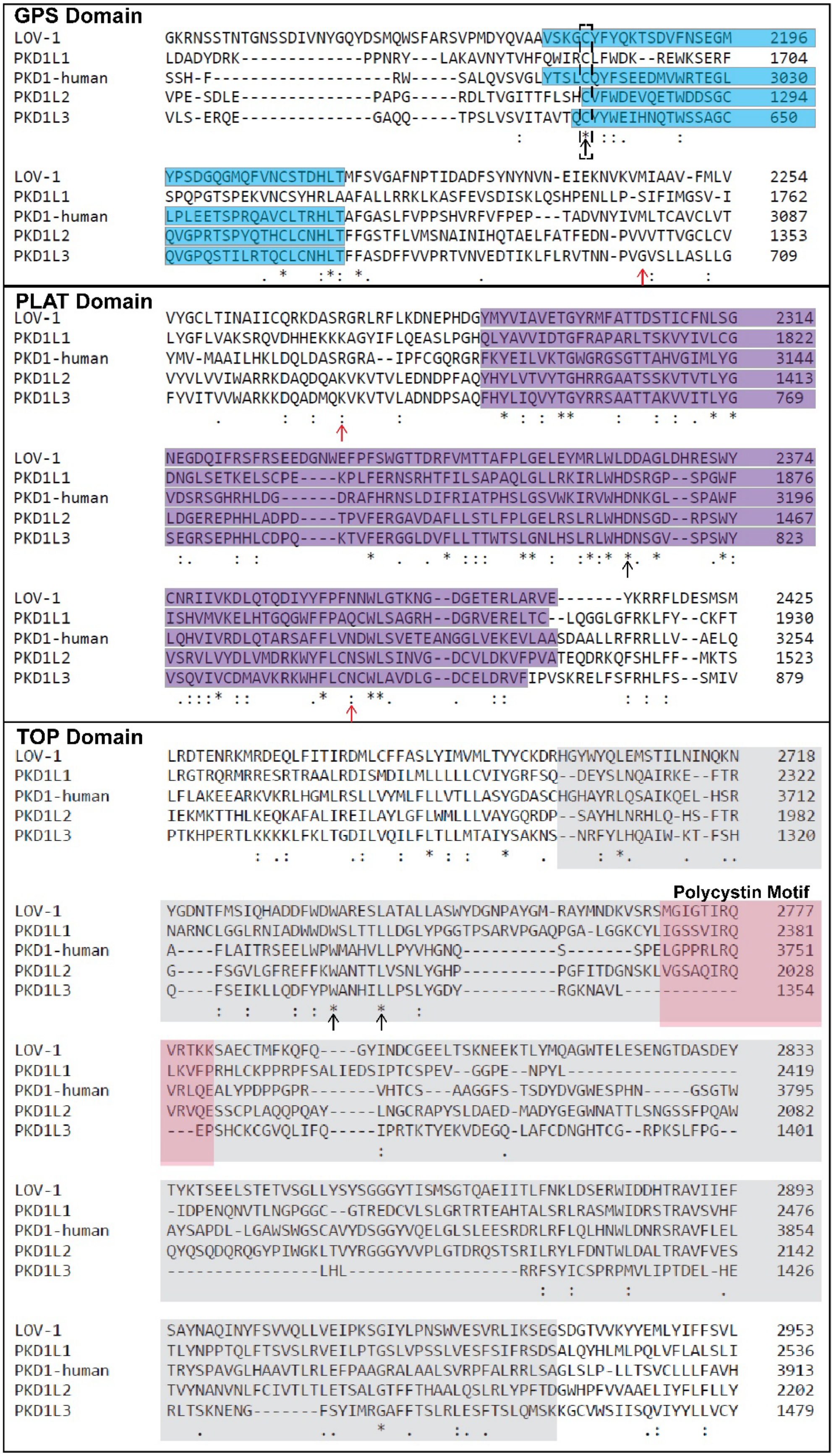
GPS Domain
LOV-1 GKRNSSTNTGNSSDIVNYGQYDSMQWSFARSVPMDYQVAAVSKGCYFYQKTSDVFNSEGM 2196
PKD1L1 LDADYDRK-------------PPNRY---LAKAVNYTVHFQWIRCLFWDK--REWKSERF 1704
PKD1-human SSH-F-------------------RW-----SALQVSVGLYTSLCQYFSEEDMVWRTEGL 3030
PKD1L2 VPE-SDLE--------------PAPG-----RDLTVGITTFLSHCVFWDEVQETWDDSGC 1294
PKD1L3 VLS-ERQE--------------GAQQ-----TPSLVSVITAVTQCVYWEIHNQTWSSAGC 650
LOV-1 YPSDGQGMQFVNCSTDHLTMFSVGAFNPTIDADFSYNYNVN-EIEKNVKVMIAAV-FMLV 2254
PKD1L1 SPQPGTSPEKVNCSYHRLAAFALLRRKLKASFEVSDISKLQSHPENLLP-SIFIMGSV-I 1762
PKD1-human LPLEETSPRQAVCLTRHLTAFGASLFVPPSHVRFVFPEP---TADVNYIVMLTCAVCLVT 3087
PKD1L2 QVGPRTSPYQTHCLCNHLTFFGSTFLVMSNAINIHQTAELFATFEDN-PVVVTTVGCLCV 1353
PKD1L3 QVGPQSTILRTQCLCNHLTFFASDFFVVPRTVNVEDTIKLFLRVTNN-PVGVSLLASLLG 709
PLAT Domain
LOV-1 VYGCLTINAIICQRKDASRGRLRFLKDNEPHDGYMYVIAVETGYRMFATTDSTICFNLSG 2314
PKD1L1 LYGFLVAKSRQVDHHEKKKAGYIFLQEASLPGHQLYAVVIDTGFRAPARLTSKVYIVLCG 1822
PKD1-human YMV-MAAILHKLDQLDASRGRA--IPFCGQRGRFKYEILVKTGWGRGSGTTAHVGIMLYG 3144
PKD1L2 VYVLVVIWARRKDAQDQAKVKVTVLEDNDPFAQYHYLVTVYTGHRRGAATSSKVTVTLYG 1413
PKD1L3 FYVITVVWARKKDQADMQKVKVTVLADNDPSAQFHYLIQVYTGYRRSAATTAKVVITLYG 769
LOV-1 NEGDQIFRSFRSEEDGNWEFPFSWGTTDRFVMTTAFPLGELEYMRLWLDDAGLDHRESWY 2374
PKD1L1 DNGLSETKELSCPE----KPLFERNSRHTFILSAPAQLGLLRKIRLWHDSRGP--SPGWF 1876
PKD1-human VDSRSGHRHLDG------DRAFHRNSLDIFRIATPHSLGSVWKIRVWHDNKGL--SPAWF 3196
PKD1L2 LDGEREPHHLADPD----TPVFERGAVDAFLLSTLFPLGELRSLRLWHDNSGD--RPSWY 1467
PKD1L3 SEGRSEPHHLCDPQ----KTVFERGGLDVFLLTTWTSLGNLHSLRLWHDNSGV--SPSWY 823
LOV-1 CNRIIVKDLQTQDIYYFPFNNWLGTKNG--DGETERLARVE-------YKRRFLDESMSM 2425
PKD1L1 ISHVMVKELHTGQGWFFPAQCWLSAGRH--DGRVERELTC--LQGGLGFRKLFY--CKFT 1930
PKD1-human LQHVIVRDLQTARSAFFLVNDWLSVETEANGGLVEKEVLAASDAALLRFRRLLV--AELQ 3254
PKD1L2 VSRVLVYDLVMDRKWYFLCNSWLSINVG--DCVLDKVFPVATEQDRKQFSHLFF--MKTS 1523
PKD1L3 VSQVIVCDMAVKRKWHFLCNCWLAVDLG--DCELDRVFIPVSKRELFSFRHLFS--SMIV 879
TOP Domain
LOV-1 LRDTENRKMRDEQLFITIRDMLCFFASLYIMVMLTYYCKDRHGYWYQLEMSTILNINQKN 2718
PKD1L1 LRGTRQRMRRESRTRAALRDISMDILMLLLLLCVIYGRFSQ--DEYSLNQAIRKE--FTR 2322
PKD1-human LFLAKEEARKVKRLHGMLRSLLVYMLFLLVTLLASYGDASCHGHAYRLQSAIKQEL-HSR 3712
PKD1L2 IEKMKTTHLKEQKAFALIREILAYLGFLWMLLLVAYGQRDP--SAYHLNRHLQ-HS-FTR 1982
PKD1L3 PTKHPERTLKKKKLFKLTGDILVQILFLTLLMTAIYSAKNS--NRFYLHQAIW-KT-FSH 1320
Polycystin Motif
LOV-1 YGDNTFMSIQHADDFWDWARESLATALLASWYDGNPAYGM-RAYMNDKVSRSMGIGTIRQ 2777
PKD1L1 NARNCLGGLRNIADWWDWSLTTLLDGLYPGGTPSARVPGAQPGA-LGGKCYLIGSSVIRQ 2381
PKD1-human A----FLAITRSEELWPWMAHVLLPYVHGNQ----------S-------SPELGPPRLRQ 3751
PKD1L2 G----FSGVLGFREFFKWANTTLVSNLYGHP----------PGFITDGNSKLVGSAQIRQ 2028
PKD1L3 Q----FSEIKLLQDFYPWANHILLPSLYGDY----------RGKNAVL------------ 1354
LOV-1 VRTKKSAECTMFKQFQ----GYINDCGEELTSKNEEKTLYMQAGWTELESENGTDASDEY 2833
PKD1L1 LKVFPRHLCKPPRPFSALIEDSIPTCSPEV--GGPE--NPYL------------------ 2419
PKD1-human VRLQEALYPDPPGPR-------VHTCS---AAGGFS-TSDYDVGWESPHN-----GSGTW 3795
PKD1L2 VRVQESSCPLAQQPQAY-----LNGCRAPYSLDAED-MADYGEGWNATTLSNGSSFPQAW 2082
PKD1L3 ---EPSHCKCGVQLIFQ-----IPRTKTYEKVDEGQ-LAFCDNGHTCG--RPKSLFPG-- 1401
LOV-1 TYKTSEELSTETVSGLLYSYSGGGYTISMSGTQAEIITLFNKLDSERWIDDHTRAVIIEF 2893
PKD1L1 -IDPENQNVTLNGPGGC--GTREDCVLSLGRTRTEAHTALSRLRASMWIDRSTRAVSVHF 2476
PKD1-human AYSAPDL-LGAWSWGSCAVYDSGGYVQELGLSLEESRDRLRFLQLHNWLDNRSRAVFLEL 3854
PKD1L2 QYQSQDQRQGYPIWGKLTVYRGGGYVVPLGTDRQSTSRILRYLFDNTWLDALTRAVFVES 2142
PKD1L3 ----------------LHL------------------RRFSYICSPRPMVLIPTDEL-HE 1426
LOV-1 SAYNAQINYFSVVQLLVEIPKSGIYLPNSWVESVRLIKSEGSDGTVVKYYEMLYIFFSVL 2953
PKD1L1 TLYNPPTQLFTSVSLRVEILPTGSLVPSSLVESFSIFRSDSALQYHLMLPQLVFLALSLI 2536
PKD1-human TRYSPAVGLHAAVTLRLEFPAAGRALAALSVRPFALRRLSAGLSLP-LLTSVCLLLFAVH 3913
PKD1L2 TVYNANVNLFCIVTLTLETSALGTFFTHAALQSLRLYPFTDGWHPFVVAAELIYFLFLLY 2202
PKD1L3 RLTSKNENG-------FSYIMRGAFFTSLRLESFTSLQMSKKGCVWSIISQVIYYLLVCY 1479

**S10 Fig.** Conserved domain alignment of the polycystin 1 family proteins in humans and *C. elegans*. GPS domain highlighted in blue, PLAT domain in magenta, TOP domain in grey, and the polycystin motif in red. Black arrows indicate conserved residues among all aligned proteins that are associated with ADPKD missense mutations in humans. Red arrows indicate residues that are only conserved among PKD1 and LOV- that are associated with ADPKD missense mutations in humans. The dashed box indicates the conserved Cysteine residue of the GPS that was mutated in this study (C2181S).

**TOP Domain**

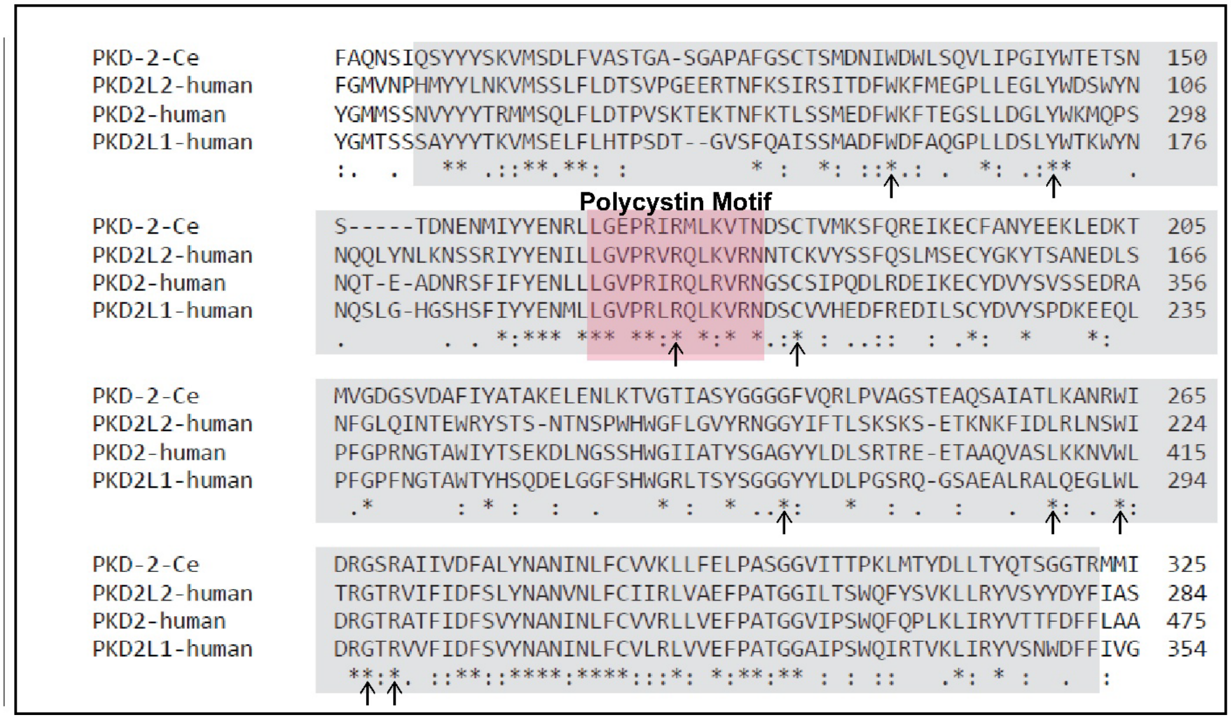

**S11 Fig.** **Conserved domain alignment of the polycystin 2 family proteins in humans and *C. elegans*.** TOP domain in grey and the polycystin motif in red. Black arrows indicate conserved residues among all aligned proteins that are associated with ADPKD missense mutations in humans.

**TOP Domain**

```
PKD1-human              HGHAYRLQSAI------KQELHSRAFLAITRSEELWPWMAHVLLPYVH------------
LOV-1C_elegans          -GYWYQLEMSTILNIN-QKNYGDNTFMSIQHADDFWDWARESLATALLASWY---DGNPA
PKD-2-C_elegans         QSYYYSKVMSDLFVAS-TGASGAPAFGSCTSMDNIWDWLSQVLIPGIY--WT---ETSNS
PKD2-human              -VYYYTRMMSQLFLDTPVSKTEKTNFKTLSSMEDFWKFTEGSLLDGLY--WKMQPSNQTE
                          : *    :               * :    :::* :    *   :
                                                           ↑

                                      Polycystin Motif
PKD1-human              -GNQS------SPELGPPRLRQVRLQE---ALYPDPPGPRVHTCSAAGGFSTSDYDVG--
LOV-1C_elegans          YGMRAYMNDKVSRSMGIGTIRQVRTKKSAECTMFKQFQGYINDCGEELTSKNEEKTLYMQ
PKD-2-C_elegans         TDNENMIYYE-NRLLGEPRIRMLKVTNDS-CTVMKSFQREIKECFANYEEKLEDKTMV--
PKD2-human              ADNRSFIFYE-NLLLGVPRIRQLRVRNGS-CSIPQDLRDEIKECYDVYSVSSEDRAPF--
                         .         .  :*   :* :.  :   .   .     :: *      . .:
                                            ↑

PKD1-human              -------WESPHNGSGTWAY-SAPDLLGAWSWGSCAVYDSGGYVQELGLS-LEESRDRLR
LOV-1C_elegans          AGWTELESENGTDASDEYTYKTSEELSTETVSGLLYSYSGGGYTISMSGT-QAEIITLFN
PKD-2-C_elegans         ----------GDGSVDAFIYATAKELENLKTVGTIASYGGGGFVQRLPVAGSTEAQSAIA
PKD2-human              ----------GPRNGTAWIYTSEKDLNGSSHWGIIATYSGAGYYLDLSRT-REETAAQVA
                                         : * :  :*      *    *...*:   :  :   *    .
                                                                 ↑

PKD1-human              FLQLHNWLDNRSRAVFLELTRYSPAVGLHAAVTLRLEFPAAGRALAALSVRPFALRRLSA
LOV-1C_elegans          KLDSERWIDDHTRAVIIEFSAYNAQINYFSVVQLLVEIPKSGIYLPNSWVESVRLIKSEG
PKD-2-C_elegans         TLKANRWIDRGSRAIIVDFALYNANINLFCVVKLLFELPASGGVITTPKLMTYDLLTYQT
PKD2-human              SLKKNVWLDRGTRATFIDFSVYNANINLFCVVRLLVEFPATGGVIPSWQFQPLKLIRYVT
                         *.   *:*  :** ::::: *.. :. ...* * .*:* :*   :.   . .  *
                         ↑    ↑     ↑                       ↑

PKD1-human              ----
LOV-1C_elegans          SDGT
PKD-2-C_elegans         ----
PKD2-human              ----
```

__S12 Fig.__ **Multiple sequence alignment of human PC1, human, PC2, *C. elegans* LOV-1, and *C. elegans* PKD-2 TOP domains.** The polycystin motif is highlighted in red. Black arrows indicate conserved residues among all aligned protein domains that are associated with ADPKD missense mutations in humans. These residues may be mutated in human PC1, PC2, or both.

**S1 Table. Strains used in this study.** All newly generated strains are available upon request.

| Strain ID | Genotype |
|---|---|
| CB1490 | him-5(e1490)V |
| PS622 | dpy-17(e164) III; him-5(e1490) V |
| CB369 | unc-51 (e369) V |
| CB444 | unc-52(e444) II |
| PS3401 | lov-1(sy582) II; him-5(e1490) V |
| PT9 | pkd-2(sy606) IV; him-5(e1490) V |
| PT443 | myIs1[PKD-2::GFP+Punc-122::GFP] pkd-2(sy606) IV; him-5(e1490) V |
| PT657 | lov-1(sy582) III; myIs4[PKD-2::GFP+Punc-122::GFP],him-5(e1490)V |
| PT2317 | myIs1[PKD-2::GFP+Punc-122::GFP]IV; him-5(e1490) V |
| PT2519 | myIs13 [Pklp-6::gfp + pBx] III; him-5(e1490) V |
| PT3424 | lov-1(my52[mSC::lov-1])II; him-5(e1490)V |
| PT3444 | lov-1(my52[mSC::lov-1])II; MyIs1[pkd-2::GFP+ccGFP]IV; him-5(e1490)V |
| PT3497 | lov-1(my54[lov-1::mNG])II; him-5(e1490)V |
| PT3498 | lov-1(my64[mSC::lov-1::mNG])II; him-5(e1490)V |
| PT3516 | lov-1(my76[mSC::lov-1[C2181S]])II;myIs1[PKD-2::GFP+Punc-122::GFP]IV; him-5(e1490)V |
| PT3517 | lov-1(my73[mNG::lov-1::mSC])II; him-5(e1490)V |
| PT3519 | lov-1(my64[mSC::lov-1::mNG])II; klp-6(my8)III; him-5(e1490)V |
| PT3530 | lov-1(my84[(my64)mSC::lov-1(C2181S)::mNG])II; him-5(e1490)V |
| PT3532 | lov-1(my64[mSC::lov-1::mNG])II; pkd-2(sy606)IV; him-5(e1490)V |
| PT3620 | lov-1(my78[lov-1::mSC])II;myIs1[PKD-2::GFP+Punc-122::GFP]IV; him-5(e1490)V |
| PT3629 | lov-1(my78[lov-1::mSC])II; him-5(e1490)V |

**Table 2:** DNA and RNA sequences used

| **crRNA homology sequence** | | color code |
|---|---|---|
| *lov-1 CTM* | GAAGTGGCGGCTAAACGATG **TGG** (Forward Strand) | **homology** |
| *lov-1 NTM* | **CCA** ACTTCTTGACGGGATCGCAA (Reverse strand) | **PAM** |
| *lov-1(C2181S)* | CAAGTCGCTGCAGTGAGTAA **AGG** (Forward Strand) | Silent mutation |
| | | Flexible Linker |
| **pdsDNA Primers** | | **mNeonGreen** |
| *lov-1::mneongreen* | | mScarlet |
| Long_Forward | CTGGACCAAAGAGATTCCAGAAGTGGCGGCTAAACGACGTCGAGAAAGATGGAGGTGGCGGATCTGGAGGTGGAGGCTCTGGAGGAGGTGGATCTATGGTGTCGAAGGGAGAAGA | |
| Long_Reverse | GAATGGATGAACTCTACAAGTAGTTTTTGAGGGCTACCTAATTTGGATATCAAGAATA | |
| Short_Forward | ATGGTGTCGAAGGGAGAAGA | |
| Short_Reverse | GAATGGATGAACTCTACAAGTAG | |
| *mscarlet-I::lov-1* | | |
| Long_Forward | GAAGGAAAAGCTGAATAAAACTACTTTTAAGATACAAATTGATGGAATGGTCAGCAAGGGAGAGG | |
| Long_Reverse | GGAATGGACGAGCTCTACAAGGGAGGTGGCGGATCTGGAGGTGGAGGCTCTGGAGGAGGTGGATCTTTGCATTAtCAgCTTCTTGACGGGATCGCAACTTTTCGATTAGACAACG | |
| Short_Forward | AAATTGATGGAATGGTCAGC | |
| Short_Reverse | GGATCTTTGCATTATCAGCT | |
| *lov-1::mscarlet-I* | | |
| Long_Forward | CTGGACCAAAGAGATTCCAGAAGTGGCGGCTAAACGAcGTcGAGAAAGATGGAGGTGGCGGATCTGGAGGTGGAGGCTCTGGAGGAGGTGGATCTATGGTCAGCAAGGGAGAGGC | |
| Long_Reverse | TATTCTTGATATCCAAATTAGGTAGCCCTCAAAAACTACTTGTAGAGCTCGTCCA | |
| Short_Forward | ACGTCGAGAAAGATGGAGGT | |
| Short_Reverse | CTACTTGTAGAGCTCGTCCA | |
| *mneongreen::lov-1* | | |
| Long_Forward | GAAGGAAAAGCTGAATAAAACTACTTTTAAGATACAAATTGATGGAATGGTGTCGAAGGGAGAAGA | |
| Long_Reverse | CGTTGTCTAATCGAAAAGTTGCGATCCCGTCAAGAAGcTGaTAATGCAAAGATCCACCTCCTCCAGAGCCTCCACCTCCAGATCCGCCACCTCCCTTGTAGAGTTCATCCATTC | |
| Short_Forward | AATTGATGGAATGGTGTCGA | |

| | |
|---|---|
| Short_Reverse | AGCTGATAATGCAAAGATCC |
| **ssODNs** | |
| *lov-1(C2181S)* | GCACGAAGTGTACCAATGGATTATCAAGTCGCTGCCGTCT GGAAAGGA **(TCC)** **TACTTCTATCAGAAAACATCGGATGTCTTCAATTCTG** **(TGT --> TCC = Cysteine --> Serine)** |
| **Genotyping Primers** | |
| *lov-1::mneongreen* | |
| Outside | Forward:CAGACAAAACGTCGCTTGGG<br>Reverse:TGACCCATCATGCCTTTGTTC |
| Inside | Forward:CAGACAAAACGTCGCTTGGG<br>Reverse:TGTGCGATCCCTCGTATGTG |
| *mscarlet-I::lov-1* | |
| Outside | Forward:CCGCCTTTTCGCTTTTCGAC<br>Reverse:CCGCCTTTTCGCTTTTCGAC |
| Inside | Forward:CCGCCTTTTCGCTTTTCGAC<br>Reverse:AGTAGTCTGGGATGTCGGCT |
| *lov-1::mscarlet-I* | |
| Outside | Forward:CAGACAAAACGTCGCTTGGG<br>Reverse:TGACCCATCATGCCTTTGTTC |
| Inside | Forward:CAGACAAAACGTCGCTTGGG<br>Reverse:AGTAGTCTGGGATGTCGGCT |
| *mneongreen::lov-1* | |
| Outside | Forward:CCGCCTTTTCGCTTTTCGAC<br>Reverse:CCGCCTTTTCGCTTTTCGAC |
| Inside | Forward:CCGCCTTTTCGCTTTTCGAC<br>Reverse:TGTGCGATCCCTCGTATGTG |
| *lov-1(C2181S)* | Forward: TATGTGAACGGCAGGGGAAG<br>Reverse:GTGGTTCCCCATGAAAACGG Restriction Digest: BamHI |
| **Gene Fusion Sequences** | |
| *lov-1::mneongreen* | **CTGGACCAAAGAGATTCCAGAAGTGGCGGCTAAACG**ACG**T** **C**GAGAAAGATGGAGGTGGCGGATCTGGAGGTGGAGGCTCTG GAGGAGGTGGATCT**ATGGTGTCGAAGGGAGAAGAGGATAA** **CATGG** |
| *mscarlet-I::lov-1* | **ACGTCACTCCACCGGAGGAATGGACGAGCTCTACAAG**GGA GGTGGCGGATCTGGAGGTGGAGGCTCTGGAGGAGGTGGATCT TTGCATTA**TCAGCTTCTTGACGGGATCGCAACTTTTCGATT** **AGACAACG** |

| | |
|---|---|
| *lov-1::mscarlet-I* | **CTGGACCAAAGAGATTCCAGAAGTGGCGGCTAAACG**ACG**TCG**AGAAAGATGGAGGTGGCGGATCTGGAGGTGGAGGCTCTGGAGGAGGTGGATCT**ATGGTCAGCAAGGGAGAGGCAGTTATCAAGGAGTTCA** |
| *mneongreen::lov-1* | **CAGATGTGATGGGAATGGATGAACTCTACAAG**GGAGGTGGCGGATCTGGAGGTGGAGGCTCTGGAGGAGGTGGATCTTTGCATTA**TCA**G**CTTCTTGACGGGATCGCAACTTTTCGATTAGACAACG** |
